# Modelling the emergence of whisker barrels

**DOI:** 10.1101/2019.12.20.884106

**Authors:** Sebastian S. James, Leah A. Krubitzer, Stuart P. Wilson

## Abstract

Brain development relies on an interplay between genetic specification and self-organization. Striking examples of this relationship can be found in the somatosensory brainstem, thalamus, and cortex of rats and mice, where the arrangement of the facial whiskers is preserved in the arrangement of cell aggregates to form precise somatotopic maps. We show in simulation how realistic whisker maps can self-organize, by assuming that information is exchanged between adjacent cells only, under the guidance of gene expression gradients. The resulting model provides a simple account of how patterns of gene expression can constrain spontaneous pattern formation to faithfully reproduce functional maps in subsequent brain structures.

## Introduction

Spatial patterns in neural connectivity provide clues about the constraints under which brains evolve and develop (***Purves et al., 1992***). Perhaps the most distinctive pattern can be found in the barrel cortex of many rodent species (***Woolsey and Van der Loos, 1970***). The barrels are identifiable soon after birth in layer 4 of primary somatosensory cortex as dense clusters of thalamocortical axons, which are enclosed by borders a few neurons thick from postnatal day 3 (***Erzurumlu and Gaspar, 2012***). In the plane tangential to the cortical surface the barrels constitute a somatotopic map of the whiskers, with cells within adjacent barrels responding most strongly and quickly to deflection of adjacent whiskers (***Armstrong-James et al., 1992***). Barrel patterning reflects subcortical whisker maps comprising cell aggregates called barrelettes in the brainstem and barreloids in the thalamus (***Ma, 1991***; ***Van Der Loos, 1976***).

Barrel formation requires afferent input from whisker stimulation and thalamic calcium waves (***Antón-Bolaños et al., 2019***), and depends on a complex network of axon guidance molecules such as ephrin-A5 and A7 and adhesion molecules such as cadherin-6 and 8 (***Vanderhaeghen et al., 2000***; ***Miller et al., 2006***). This network is orchestrated by interactions between morphogens Fgf8 and Fgf17 and transcription factors Emx2, Pax6, Sp8, and Coup-tf1 (***Shimogori and Grove, 2005***; ***Bishop et al., 2000***), which are expressed in gradients that mark orthogonal axes and can be manipulated to stretch, shrink, shift, and even duplicate barrels (***Assimacopoulos et al., 2012***).

The barrel boundaries form a Voronoi tessellation across the cortical sheet (***Senft and Woolsey, 1991***) (Fig. 1A), suggesting that barreloid topology is preserved in the projection of thalamocortical axons into the cortex, and that a barrel forms by lateral axon branching from an initial center-point that ceases upon contact with axons branching from adjacent centers. However, the assumption of pre-arranged center-points is difficult to resolve with the observation that axons arrive in the cortical plate as an undifferentiated bundle, *prior* to barreloid formation (***Agmon et al., 1993***).

**Figure 1.**
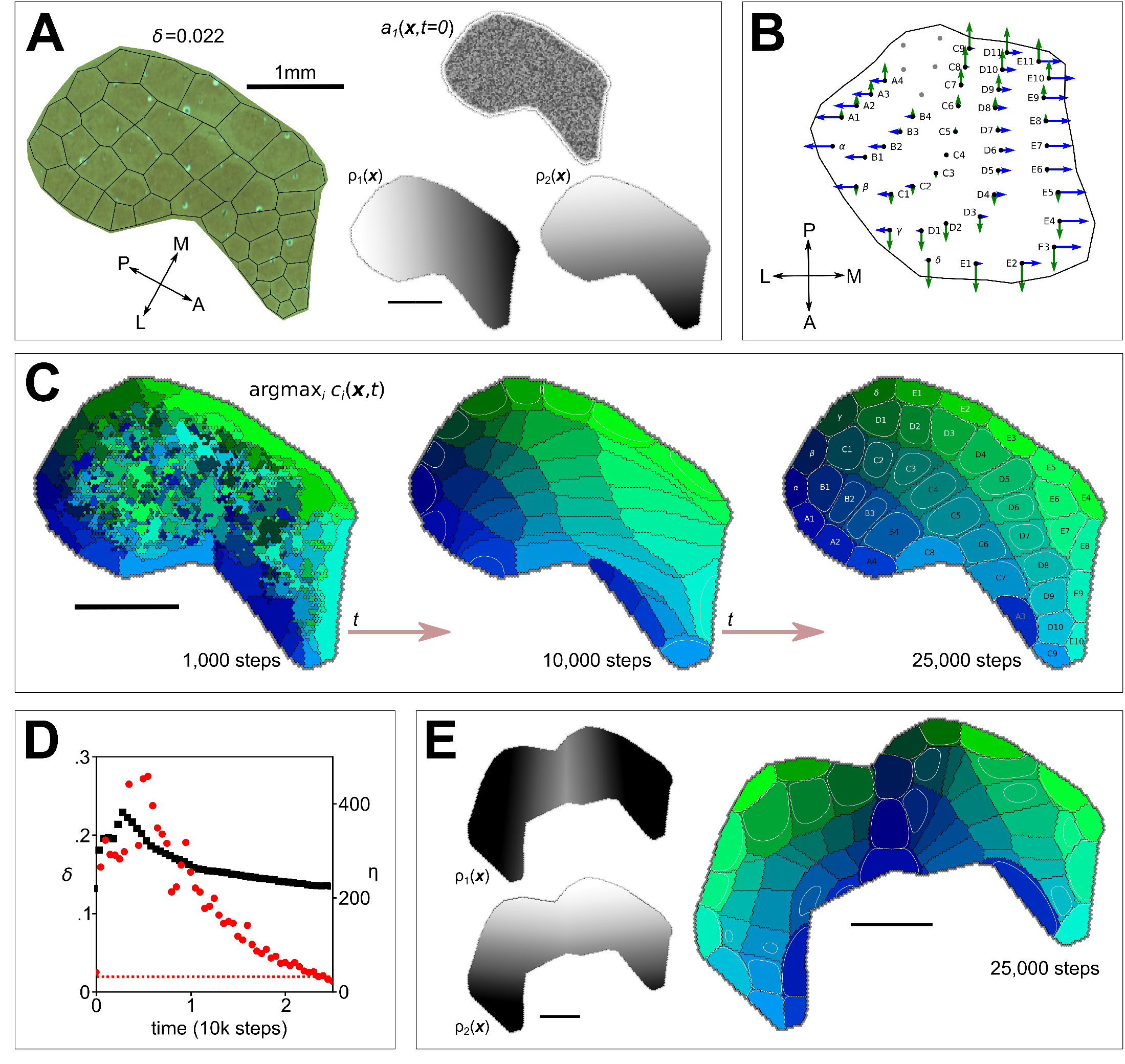
**A** Left shows a cytochrome oxidase stain obtained from rat S1 by ***Zheng et al. (2001)***, with black lines to delineate barrels and to measure departure (Honda-*δ*; see ***Senft and Woolsey, 1991***) from a perfect Voronoi tesselation. Right shows the initial distribution of axon branching density (*a*) for one thalamocortical projection, and two molecular guidance fields (*ρ*). **B** The strengths of interaction *γ* with fields *ρ*_1_ and *ρ*_2_ are indicated for each of 41 projections by the lengths of green and blue arrows respectively, assuming that similar fields aligned to the posterior-anterior and medial-lateral axes in the ventroposterior medial nucleus of the thalamus are sampled at the locations of putative barreloid centers (reconstructed from ***Haidarliu and Ahissar, 2001***). **C** Simulation results for parameters *N* = 41, *α* = 3, *β* = 20, *k* = 3, *D* = 0.2, *γ* ∈ ±2, *ϵ* = 150 and *δt* = 0.0001. Colours indicate the thalamic projection for which the connection density is maximal, black lines delineate boundaries, and overlaid contours show *c* > 0.5 (see Movie S1). **D** Red dots show the Honda-*δ* metric obtained from simulation approaching that obtained from barrels in A (dotted line); black squares measure the correspondence between the real and simulated barrel shapes, *η* (the product of the sum of squared differences between real and simulated centers and the sum of differences in area; units mm^4^). **E** Guidance fields and emergent barrel pattern in a Fgf8 misexpression experiment (c.f. ***Assimacopoulos et al., 2012***), simulated by reflecting *ρ*_1_ at the join (*ϵ* = 80). All scale bars 1mm.

Alternatively, reaction-diffusion dynamics could generate a Voronoi tessellation without pre-arranged centers, by amplifying characteristic modes in a noisy initial distribution of axon branches, as a net effect of short-range cooperative and longer-range competitive interactions. Accordingly, the barrel pattern would be determined by the relative strength of these interactions and by the shape of the cortical field boundary. However, intrinsic cortical dynamics alone cannot account for the topographic correspondence between thalamic and cortical domains, the irregular sizes and specific arrangement of the barrels in rows and arcs, or the influence of gene expression gradients.

The center-point and reaction-diffusion models are not mutually exclusive. Pre-organized centers could bias reaction-diffusion processes to generate specific arrangements more reliably, and mechanisms of lateral axon branching may constitute the tension between cooperation and competition required for self-organization. However, proof that barrel patterning can emerge from an undifferentiated bundle of axons, based only on local interactions, would show that a separate stage and/or extrinsic mechanism for pre-organizing thalamocortical connections need not be assumed. To this end, we ask whether barrel maps can emerge in a system with reaction-diffusion dynamics, under the guidance of signalling gradients, and in the absence of pre-defined centers.

## Models

***Karbowski and Ermentrout(2004)*** developed a reaction-diffusion style model of how extrinsic signalling gradients can constrain the emergence of distinct fields from intrinsic cortical dynamics. Their model defines how the density of connections *c*(*x,t*) and axon branches *a*(*x,t*) interact at time *t*, along a 1D anterior-posterior axis *x*, for *N* thalamocortical projections indexed by *i*. The model was derived from the assumption that the rates at which *a*_*i*_ and *c*_*i*_ grow are reciprocally coupled. Extending the original 1D model to simulate arealization on a 2D cortical sheet, we use *a*_*i*_(**x**,*t*) and *c*_*i*_(**x**,*t*), and model synaptogenesis as

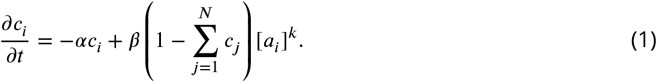

Accordingly, where the total density of synaptic connections sums to one, connections decay at rate *α*. Otherwise the connection density increases non-linearly (*k* > 1) with the density of axon branching. Axon branching is modelled as

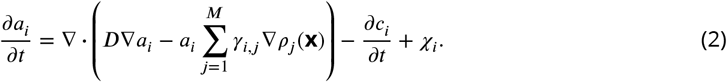

The first term on the right describes the divergence (indicated by ∇·) of the quantity in parentheses, which is referred to as the ‘flux’ of axonal branching. The flux represents diffusion across the cortical sheet, at rate *D*, and the influence of *M* molecular signalling fields, *ρ*(**x**). The influence of a given field (indexed by *j*) on a given thalamic projection (indexed by *i*), is determined by *γ*_*i,j*_, which may be positive or negative in order that axons may branch in the direction of either higher or lower concentrations. Note that computing the divergence in simulation requires cells on the cortical sheet to communicate with immedately adjacent cells only (see *Materials & Methods*) The second term on the right quantifies the coupling between axon branching and synaptogenesis. Here *χ*_*i*_ = 0 is a placeholder.

## Results

First we verified that all results established by ***Karbowski and Ermentrout (2004)*** for a 1D axis could be reproduced using our extension to a 2D cortical sheet. Using an elliptical domain, *S*, with *M* = 3 offset guidance gradients aligned to the longer axis, *N* = 5 thalamocortical projections gave rise to five distinct cortical fields at locations that preserved the topographic ordering defined by the original *γ* values. However, we found that specifying *N* ordered areas required *M* ≈ (*N* + 1)/2 signalling fields. This is because localization of axon densities occurs only when projections are influenced by interactions with two or more signalling gradients that encourage migration in opposing directions.

Note that these dynamics are quite unlike classic chemospecificity models (***Sperry, 1963***), which essentially assume center-points i.e., conditions in the target tissue that instruct pre-identified afferents to stop growing. As the number of distinct guidance fields is unlikely to approach the number of barrels, further extensions were required.

The term in parentheses in Eq. 1 represents competition between thalamocortical projections for a limited availability of cortical connections. To introduce competition also in terms of axon branching we redefined

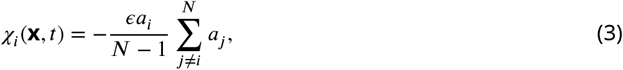

which reduces branching for each projection where the branches of other projections are dense. Divisive normalization keeps the axon branch density bounded on each iteration to the initial (random) density of unconnected branches at *t* = 0

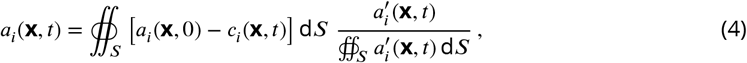

where prime symbols indicate the use of intermediate values computed from Eqs. 1–3 at time *t* − *δt* (*δt* is the time represented by one simulation step). Note that this operation is local to individual afferent projections.

The only differences between thalamic projections are their strengths of interaction with the guidance fields, *γ*, hence any reliable differences between emergent cortical fields must be due to differences in these values only. We speculate that the contribution of a given ephrin field, to the velocity at which a projection migrates across the cortical subplate, is determined by the concentration of a similar molecule at its thalamic origin, i.e., the putative barreloid center. As such, two orthogonal linear thalamic gradients were defined, from which 41 pairs of values were sampled, at the coordinates of 41 barreloid centers recreated from ***Haidarliu and Ahissar (2001)*** (Fig. 1B).

A cortical boundary enclosing 41 corresponding barrels was traced from a cytochrome oxidase stain from ***Zheng et al. (2001)***, and Eqs. 1–4 were solved for *N* = 41 projections on the resulting domain, using *M* = 2 linear signalling gradients aligned with the anterior-posterior and medial-lateral axes. From random initial conditions for *a*(**x**,0) and *c*(**x**,0), a clear Voronoi-like tesselation emerged (Fig. 1C; see Movie S1). A reduction in the Honda delta metric (see ***Senft and Woolsey, 1991***) confirmed that a ‘good’ Voronoi pattern emerged within ≈ 3000 iterations (Fig. 1D). A measure of the difference in shape between real and simulated barrels revealed a strong correspondence (see Fig. 1D).

To further investigate the interplay of genetic and intrinsic mechanisms we simulated a seminal barrel duplication experiment (***Shimogori and Grove, 2005***), first by solving on a cortical domain comprising two separate, identically shaped cortical boundaries. In this case one distinct cortical field per thalamic projection emerged, within one of the two boundaries only, i.e., no duplication. However, merging the two fields to create an extended mirror-symmetric boundary shape, with an anterior-posterior guidance field reflected at the join, gave rise to two mirror-symmetrical barrel fields comprising 2N barrels, i.e., two cortical fields for each thalamic projection (Fig. 1E). Together these results suggest that by the misexpression of Fgf8, ***Shimogori and Grove (2005)*** effectively created one large barrel field rather than two distinct cortical fields.

## Discussion

The present results suggest that the key requirements for the emergence of realistic barrel patterning are i) at each cortical location thalamocortical projections compete for a limited number of available synaptic connections (Eqs. 1–2), ii) at each location the branching rate of a given projection is reduced by the density of other projections (Eq. 3), and iii) the branch density of each projection is conserved over time (Eq. 4).

The emergence of barrels in simulation required competition between thalamic projections in terms of synaptic connectivity and also competition in terms of cortical space, as represented by *χ*, with an implicit requirement for a self/other identifier amoungst projections. This latter form of competition may account for the absence of barrels in rodents with larger brains, such as capybara, for which competition for space is presumably weaker (***Woolsey et al., 1975***). Hence, irrespective of whether barrels are necessary for adaptive whisker function, the emergence of somatotopically ordered modular structures may be an inevitable consequence of local competition for cortical territory driven by input from an array of discrete sensory organs (***Purves et al., 1992***).

It is important to emphasize that the formulation of the model is entirely local, insofar as simulation requires no information to be communicated from a given cortical grid cell to any but those immediately adjacent (via diffusion). Hence the simulations demonstrate how a self-organizing system, constrained by genetically specified guidance cues and by the shape of the cortical field boundary, can faithfully reproduce an arrangement of cell aggregates in one neural structure as a topographic map in another.

Moreover, the present results confirm that somatotopic map formation does not require the pre-specification of center-points by as yet undetermined additional developmental mechanisms.

## Materials & Methods

The cortical sheet was modelled as a two dimensional hexagonal lattice, which simplifies the computation of the 2D Laplacian. Within a boundary traced around the edge of a rat barrel field (Fig. 1A) we set the hex-to-hex distance *d* to 0.03 mm, which resulted in a lattice containing 6515 hexes for the simulations shown in Figs. 1A,C & D and 12739 hexes for the Fgf8 misexpression study shown in Fig. 1E. Each hex contained 82 time-dependent variables: 41 branching densities (*a*_*i*_) and 41 connection densities (*c*_*i*_). The rate of change of each of the time-dependent variables (Eqs. 1 & 2) was computed using a fourth-order Runge-Kutta method.

The most involved part of this computation is to find the divergence of the flux of axonal branching, **J**_*i*_(**x**,*t*) the term in parentheses in Eq. 2:

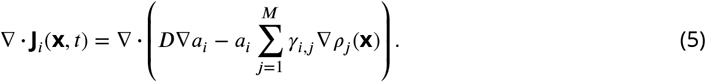

Note that the sum of the guidance gradients is time-independent and define 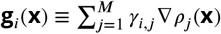. Because the divergence operator is distributive, Eq. 5 can be expanded using vector calculus identites (dropping references to **x** and *t* for clarity):

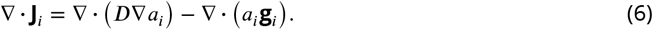

Applying the vector calculus product rule identity yields

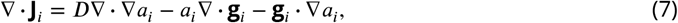

which has three elements to compute: i) *D*∇·∇*a*_*i*_ (the Laplacian of *a*_*i*_); ii) a time-independent modulator of *a*_*i*_ (because ∇·**g**_*i*_ is a time-independent static field); and iii) the scalar product of the static vector field **g**_*i*_ and the gradient of *a*_*i*_. Each of the divergences can be simplified by means of Gauss’s Theorem following ***Lee et al. (2014)***.

i. The computation of the mean value of the Laplacian across one hexagon of area 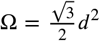, located at position **p**_0_, with neighbours at positions **p**_1_–**p**_6_ is

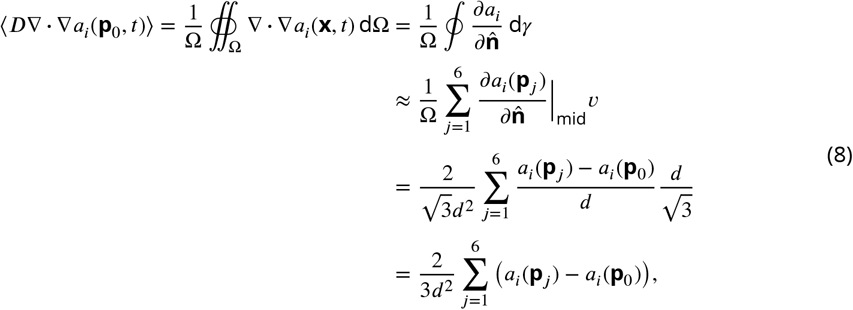

where 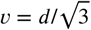 is the length of each edge of the hexagon and d*γ* is an infinitesimally small distance along its perimeter.
ii. The computation of the second term in Eq. 7, ⟨*a*_*i*_(**p**_0_,*t*)∇·**g**_*i*_(**p**_0_)⟩, can be written out similarly:

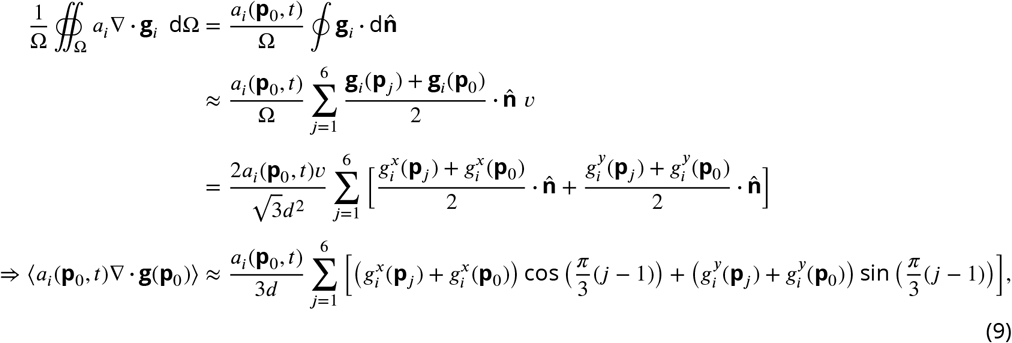

where 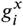 and 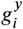 are the Cartesian components of **g**_*i*_. Both this last expression, and the final ex-168 pression of Eq. 8 can be computed locally, by summing over values of the nearest neighbours.
iii. The final term in Eq. 7 is the scalar product of two vector fields which is straightforward to compute from their Cartesian components.

By separating the computation of Eq. 5 into parts (i), (ii) & (iii), the no-flux boundary condition, **J**_*i*_(**x**,*t*)|_boundary_ = 0, can be fulfilled. On the boundary, the contribution to **J** resulting from the first term of Eq. 7 can be fixed to 0 by the ‘ghost cell method’ in which, during the evaluation of (i), a hex outside the boundary containing the same value as the hex inside the boundary is imagined to exist such that the flux of **J** across the boundary is 0. Then, **g**_*i*_(**x**) can be tailored so that it, and its normal derivative approach 0 at the boundary, ensuring that the second and third terms of Eq. 7 also contribute nothing to **J**. This is achieved by applying to **g**_*i*_(**x**) a sharp logistic function of the distance from **x** to the boundary.

All code required to reproduce these results is available at https://github.com/ABRG-Models/BarrelEmerge/tree/eLife_submission1. The computations described in (i), (ii) and (iii) may be found in the class method RD_James::compute_divJ() which calculates term1, term2 and term3, respectively.

## Movie S1 caption

Movie corresponding to Fig. 1C in the main paper. Simulation parameters were *N* = 41, *α* = 3, *β* = 20, *k* = 3, *D* = 0.2, *γ* ∈ ±2, *ϵ* = 150 and *δt* = 0.0001. Colours indicate the thalamic projection for which the connection density is maximal, black lines delineate boundaries, and overlaid contours show *c* > 0.5. The final frame in the movie is step 25,000 of the simulation.

## Acknowledgments

The authors thank Jason Berwick at the University of Sheffield for advice and for access to the rat barrel stains used to construct Fig. 1A. This work was supported by a Collaborative Activity Award, *Cortical Plasticity Within and Across Lifetimes*, from the James S. McDonnell Foundation (grant 220020516).

